# A mouse model with high clonal barcode diversity for joint lineage, transcriptomic, and epigenomic profiling in single cells

**DOI:** 10.1101/2023.01.29.526062

**Authors:** Li Li, Sarah Bowling, Qi Yu, Sean E. McGeary, Karel Alcedo, Bianca Lemke, Mark Ferreira, Allon M. Klein, Shou-Wen Wang, Fernando D. Camargo

**Author notes:** Correspondence (S.-W.W.), (F.D.C.).

## Abstract

Cellular lineage histories along with their molecular states encode fundamental principles of tissue development and homeostasis. Current lineage-recording mouse models have limited barcode diversity and poor single-cell lineage coverage, thus precluding their use in tissues composed of millions of cells. Here, we developed DARLIN, an improved Cas9 barcoding mouse line that utilizes terminal deoxynucleotidyl transferase (TdT) to enhance insertion events over 30 CRISPR target sites, stably integrated into 3 distinct genomic loci. DARLIN is inducible, has an estimated ~10^18^ lineage barcodes across tissues, and enables detection of usable barcodes in ~60% of profiled single cells. Using DARLIN, we examined fate priming within developing hematopoietic stem cells (HSCs) and revealed unique features of HSC migration. Additionally, we adapted a method to jointly profile DNA methylation, chromatin accessibility, gene expression, and lineage information in single cells. DARLIN will enable widespread high-resolution study of lineage relationships and their molecular signatures in diverse tissues and physiological contexts.

## Introduction

Tracing cellular lineage history in animals has been a long-standing effort. Historically, labeling cells with distinguishable and heritable markers such as dyes has led to major discoveries in early development and stem cell differentiation^1–3^. However, this approach is limited to tracking only small or pre-defined populations of cells. Retrovirally barcoding cells with synthetic DNA sequences has enabled analysis of much larger populations^4,5^, although this requires *ex vivo* manipulation of cells, which might alter their behavior. *In vivo* DNA barcoding in mouse models has been achieved through the use of randomly integrated transposons or recombinases that create genetic diversity within a distinct locus, which revealed a drastically different picture of hematopoiesis *in vivo*^6–10^. However, these mouse models either have limited barcode diversity, or do not allow simultaneous interrogation of lineage and state information in single cells.

The advent of CRISPR-Cas9 technology has created a new avenue for lineage tracing where diverse DNA mutations can be created within a defined locus through genome editing^11^. The mutational outcomes can be transcribed, thereby allowing joint measurement of lineage and transcriptomic information in single cells^12–14^. In mice, these approaches have been used to study early embryo development^15,16^ and cancer progression^17,18^. Applying the same tools, we developed Cas9/CARLIN, a stable and genetically defined mouse line, which enables flexible induction at any point to generate diverse, transcribed lineage barcodes across tissues^19^.

These and other single-cell lineage tracing approaches have generally faced three technical challenges: 1) low lineage barcode capture in single-cell readout; 2) low efficiency of introducing lineage barcodes (achieved by lineage barcode editing in Cas9/CARLIN); and 3) contamination from barcode homoplasy, where an identical editing event occurs independently in two different cells. As a result of these challenges, only ~10% of profiled cells from the Cas9/CARLIN mouse contain detected lineage barcodes that likely label individual clones^19^. Therefore, a higher-performing lineage-tracing mouse line is needed to enable high-coverage single-cell lineage tracing in adult tissues with millions of cells.

Single-cell lineage tracing with transcriptomic-state measurement has been successfully used to identify early fate bias among progenitors and find novel regulators of cell-fate choices^20–22^. However, epigenomic modalities such as chromatin accessibility and DNA methylation are known to play a crucial role in regulating gene expression and maintaining cell identities^23–25^. Epigenetic changes are known to foreshadow changes in gene expression^26–29^, suggesting that the earliest events for cell fate choice are unlikely to be captured using gene expression alone. This view is supported by a recent state–fate lineage tracing study showing that the transcriptome of a cell alone is insufficient to predict its fate outcome^21^. To fully understand cell fate choices and the maintenance of cell identities, single-cell approaches that integrate lineage, transcriptomic, and epigenomic information will be necessary. To this end, several recent studies have reported single-cell measurement of either lineage and transcriptomic information, lineage and epigenomic information, or transcriptomic and epigenomic information^25,30–32^. Although some multi-omic studies have inferred lineage information using endogenous DNA mutations^33,34^, the inferred clones are low-resolution and cannot be used to study lineage relationships at defined developmental time points. Indeed, a method capable of simultaneously profiling engineered lineage barcodes, the transcriptome, and the epigenome in single cells has not been reported.

Here, we developed an improved inducible and expressible lineage tracing mouse line (DARLIN) that has an extremely large lineage barcode capacity and highly efficient lineage recovery in single-cell assays, greatly outperforming Cas9/CARLIN mouse line. Furthermore, we developed a new method, Camellia-seq, to simultaneously measure DNA methylation, chromatin accessibility, gene expression, and lineage information in single cells. We applied the DARLIN mouse line and Camellia-seq method to address three distinct lineage-related problems.

## Results

### Cas9–TdT introduces more insertions than Cas9 upon transient induction in CARLIN mice

CRISPR/Cas9-based DNA editing is prone to deletions, which limits the resulting barcode diversity. Reanalyzing the editing events observed among the 10 tandem target sites within the integrated locus (referred to as target array) from the published Cas9/CARLIN mouse dataset^19^ (Figure 1A), we found that deletion events occur more frequently than insertions (Figure 1B), with 1.5 insertion events and 2.5 deletion events per allele on average. An allele from Cas9 editing has a median of 163 bp deleted out of 270 bp unedited target array, implying the deletion of 6 out of 10 tandem target sites (Figure 1C). In contrast, an allele has only a median of 2 bp inserted (Figure 1D). These large deletion events can lead to information loss and generate degenerate alleles. Indeed, alleles with only deletions lead to more common alleles, making it possible that cells with different ancestries share the same allele or lineage barcode. On the other hand, rare alleles, which label bona fide clones, overwhelmingly result from DNA insertions (Figure 1E).

**Figure 1.**
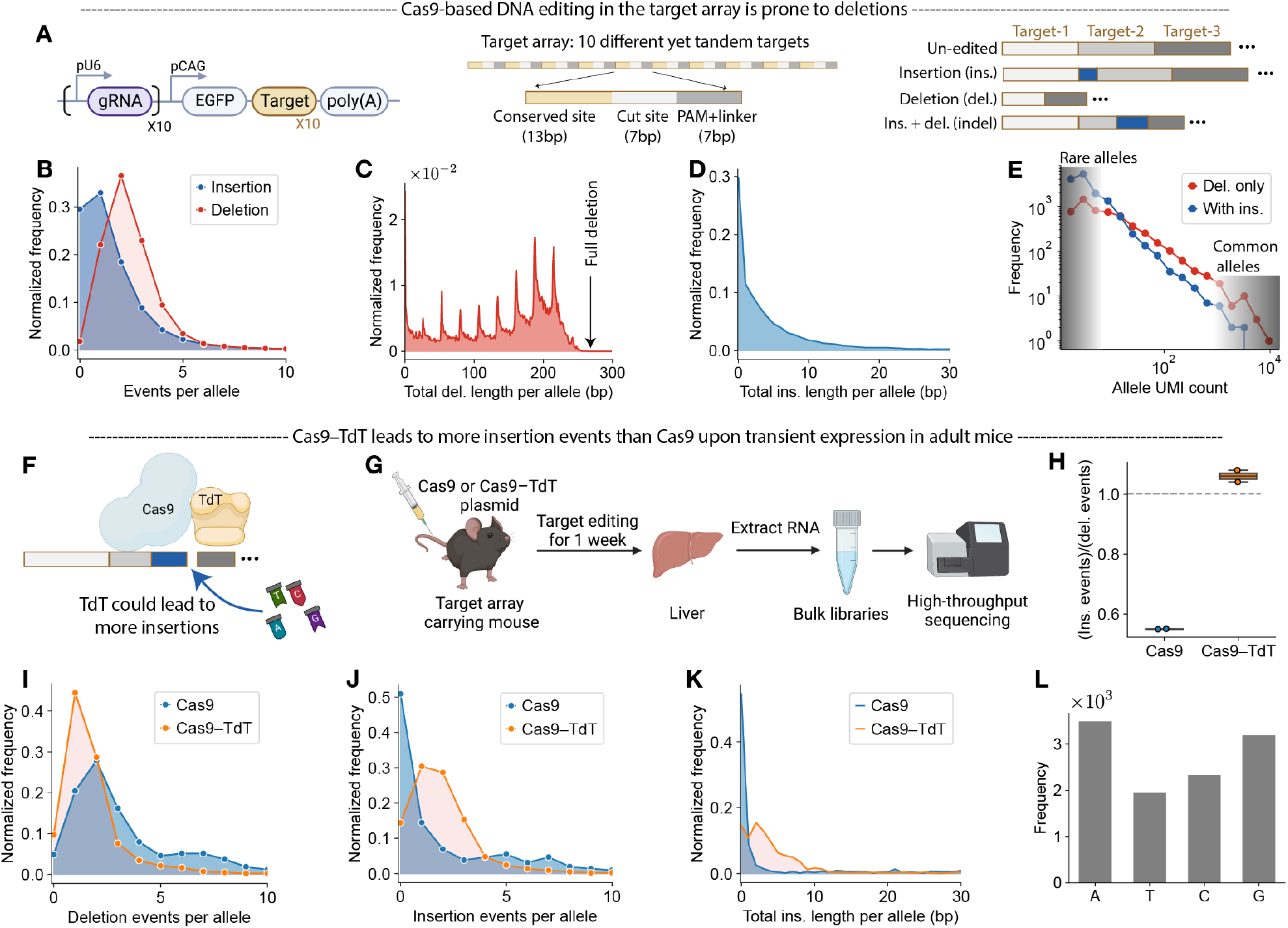
Advantage of Cas9–TdT over Cas9 for Generating Lineage Barcodes. (A) Schematic of CARLIN lineage-recording system (left), the structure of target array (middle), and several possible editing patterns within the first three target sites of the array (right). (B–E) Reanalysis of published bulk granulocyte Cas9/CARLIN data^19^. (B) Distribution of insertion or deletion events per edited allele. (C & D) Distribution of total deletion length (C) or total insertion length (D) per edited allele. (E) Histogram of allele UMI counts among edited alleles which either only contain deletions or have insertions and possibly deletions. (F) Schematic demonstrating DNA target site editing by a Cas9–TdT fusion protein. The TdT enzyme was included to increase insertion frequency. (G) Experimental scheme by which the target array editing generated from Cas9 or Cas9–TdT protein was compared. (H) Box plot showing the ratio of total insertion events to total deletion events among all edited alleles generated by editing with Cas9 or Cas9–TdT. Each point is a mouse replicate. Here and throughout, horizontal lines of each box represent the minimum, 25th-percentile, 50th-percentile, 75th-percentile, and maximum values. (I–K) Distribution of deletion (I) or insertion (J) events per allele, and total insertion length per allele (K). (L) Frequency of all four DNA nucleotides among all inserted sequences identified in the edited alleles generated by Cas9–TdT. All alleles were weighted equally.

We reasoned that increasing the frequency of insertions could greatly improve the generation of rare alleles. Terminal deoxynucleotidyl transferase (TdT) is a template-independent DNA polymerase that can insert random nucleotides at both overhang and blunt 3’ ends^35,36^. A recent study in cell lines showed that co-expression of Cas9 and TdT generated more insertions in the target sites and led to higher barcode diversity compared with Cas9 alone (Figure 1F)^37^. To test whether a similar strategy also works in an organismal context, we hydrodynamically injected a plasmid encoding either a Cas9–TdT fusion protein or native Cas9 into the tail veins of adult mice carrying the target array (Figure 1A) and analyzed the resulting allele editing in mouse livers with bulk RNA sequencing after one week (Figure 1G). We observed that Cas9–TdT expression resulted in fewer deletions but twice the insertion events per allele than Cas9 expression (Figures 1H–1J). Aggregating insertion events from all alleles, we also observed more inserted nucleotides per allele upon Cas9–TdT expression (Figure 1K), with all four nucleotides well-represented in the inserted sequences (Figure 1L). These data demonstrate that Cas9–TdT introduces more insertions as well as fewer deletions than Cas9 in the target array in an adult mouse.

### DARLIN: an inducible Cas9–TdT mouse line for high-capacity lineage tracing

Having established that the Cas9–TdT expression leads to improved lineage barcode editing in an adult mouse, we set out to generate an inducible germline mouse model that utilizes Cas9–TdT for target-array editing. First, we created a mouse embryonic stem cell (mESC) line with a Dox inducible Cas9–TdT construct also carrying a CRISPR target array and cognate gRNAs (Figure S1A). We validated that editing of the array in this mESC line was sensitive to Dox exposure: there was little editing in the absence of Dox, and higher Dox concentration or longer exposure led to more allele editing (Figures S1A-S1D).

We next engineered knock-in mice with the tetO-Cas9–TdT construct inserted into the *Cola1a* locus^38^. This line was bred with animals containing the gRNAs and *Cola1a1* target array (CA) previously described in the CARLIN system to generate *Col1a1*^tetO-Cas9–TdT/gRNA-Array^:*Rosa26*^M2-rtTA/+^ mice for lineage barcoding. We will refer to this particular line as DARLIN-v0 (Cas9-T<u>d</u>T C<u>ARLIN</u> version 0; Figure 2A). To benchmark DARLIN-v0 against the original Cas9/CARLIN mouse line, we compared their allele editing observed in large numbers of granulocytes from each mouse line after one week of Dox treatment, followed by another 3 days without Dox (Figure 2B; *n* = 7 mouse replicates for DARLIN-v0 and *n* = 5 for Cas9/CARLIN). Compared with Cas9/CARLIN, edited alleles from the DARLIN-v0 mouse were highly enriched in rare alleles (Figures 2C and S1E). Indeed, for mouse replicates with over 10,000 alleles, ~65% of alleles were observed only once (referred to as singleton alleles) for DARLIN-v0, compared with ~30% for Cas9/CARLIN (Figure 2D). For the same number of edited cells, the DARLIN-v0 mice exhibited 2.3-fold as many alleles as Cas9/CARLIN (Figure S1F). Since the utility of an allele for clonal labeling depends on its occurrence frequency, we used the metric *2^H^*, where *H* is the Shannon entropy of the normalized allele frequency across edited cells, to report the barcode diversity of an ensemble of alleles^19^. The Shannon allele diversity of DARLIN-v0 alleles was ~5 times that of Cas9/CARLIN alleles (Figure 2E). The increased diversity of the alleles from DARLIN-v0 mice is explained predominantly by the drastic reduction in the size of deletion events and also increased insertion length (Figures 2F, 2G, and S1G). Considering all mutation events across alleles, each insertion in Cas9/CARLIN was on average accompanied by three deletions, while in DARLIN-v0 each deletion implied one or more insertions (Figure 2H). There were also more insertions across the 10 target sites in DARLIN-v0 mice (Figures 2I and S1J), with all four nucleotide identities well-represented within these inserted sequences (Figure S1H). These data demonstrate that, compared with Cas9/CARLIN, the DARLIN-v0 mouse line achieves a higher fraction of rare alleles and higher barcode diversity due to more insertions and fewer deletions enabled by the Cas9–TdT editing system.

**Figure 2.**
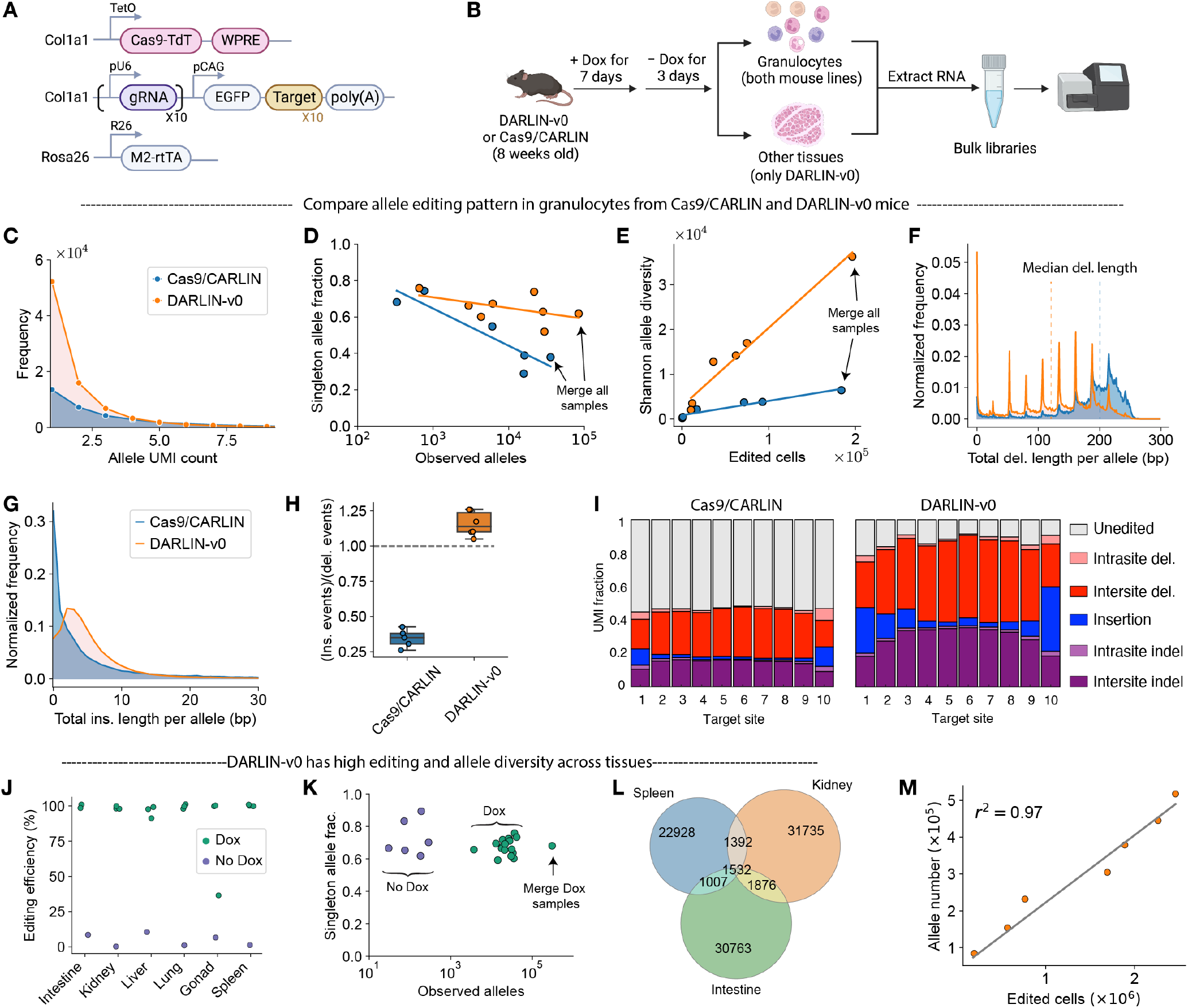
Characterization of DARLIN-v0 mice. (A) Schematics of DARLIN-v0 system. A Cas9–TdT expression cassette with a woodchuck hepatitis virus posttranscriptional regulatory element (WPRE) in its 3’ UTR is under the control of a tetO promoter at the *Col1a1* locus. The mutant tetracycline reverse transactivator (M2-rtTA) expressed from the *Rosa26* locus activates Cas9–TdT-WPRE expression upon Dox administration, leading to editing in the target array. (B) Experimental scheme to compare the target array editing in different tissues between the DARLIN-v0 and Cas9/CARLIN mice. (C–I) Comparison of the alleles generated in granulocytes of Cas9/CARLIN mice (blue) with those of DARLIN-v0 mice (orange). (C) Histogram of UMIs per allele. (D) Singleton-allele (i.e., having only one UMI) fraction as a function of the total number of observed alleles within each sample. Each point represents an individual mouse replicate except for the rightmost points, which are obtained by merging all mouse replicates within the same mouse line. (E) Shannon allele diversity as a function of edited cells approximated by total UMI counts in each sample. (F, G) Distribution of total deletion length (F) or total insertion length (G) per edited allele. (H) Box plot showing the ratio of total insertion events to total deletion events from all alleles in a sample. (I) Relative UMI fraction of editing patterns across ten target sites among alleles generated in each mouse line. (J–L) Evaluation of target-array editing in multiple tissues from the DARLIN-v0 mice. (J) Observed editing efficiency. (K) Singleton-allele fraction. (L) Venn diagram of the allele overlap between three tissues having the largest allele numbers. (M) The number of observed alleles as a function of edited cells in the DARLIN-v0 mouse line. These points correspond to merged samples from different numbers of tissue or granulocyte replicates.

### DARLIN-v0 mouse line works across tissues and yields >1 million alleles

We next sought to demonstrate that the DARLIN-v0 mouse can work across tissues, as Cas9/CARLIN did^19^. We collected intestine, kidney, lung, liver, spleen, and gonad from 3 Dox-induced DARLIN-v0 mice with the same Dox treatment as for granulocytes (Figure 2B) and 1 uninduced control DARLIN-v0 mouse. We examined the editing patterns of the target array in these tissues via bulk RNA sequencing. The editing efficiency was >90% across these tissues as compared to 30–50% in Cas9/CARLIN^19^, with only ~4% background editing without Dox induction (Figure 2J). The singleton-allele fraction was ~70%, and was comparable across tissues over a broad range of observed allele numbers (Figure 2K). Pooling alleles across all tissue samples gave a similar singleton-allele fraction, suggesting that individual tissue samples were dominated by distinct alleles. Indeed, within the same mouse, only 5% lineage barcodes were shared across the spleen, kidney, and intestine (~30,000 alleles each), indicating that most alleles were relatively rare (Figure 2L).

We next estimated the clonal-barcode diversity of DARLIN-v0 model. By progressively pooling alleles from an increasing number of mouse replicates, we observed that the number of distinct alleles increased linearly with the number of edited cells (goodness of linear fit *r*^2^ = 0.97; Figure 2M), suggesting little-to-no saturation. Pooling alleles across all available data from the DARLIN-v0 mouse line, we observed 5.2 × 10^5^ unique alleles in total (Figure 2M), with a singleton fraction of 0.62 (Figure S1I). The large fraction of singletons implies many more unobserved alleles that would be detected if we sample more cells^39^. Although our sampling was far from saturation, making it hard to infer the maximum allele number that could be produced by the DARLIN-v0 mouse, we estimated the lower bound of the total alleles to be 5.2 × 10^5^ / (1 - 0.62) = 1.3 × 10^6^ using the Good–Turing estimator^39,40^. This is at least 30 times as many as the reported 44,000 total alleles in Cas9/CARLIN mice^19^. To conclude, our DARLIN-v0 mouse line can be used for lineage tracing in various tissues and has a large barcode capacity.

### DARLIN mice contain three independant target arrays

To further increase the clonal-barcode diversity for organism-wide lineage tracing, we generated two additional mouse lines each with one target array: one integrated at the *Tigre* locus (*Tigre* target array, or TA) and the other at the *Rosa26* locus (*Rosa26* target array, or RA). TA and RA reused the same 10 target sequences from CA, so that they can be edited by the existing 10 tandem gRNAs, but in different orders (Figure S2A). We generated an additional tetO-Cas9–TdT knock-in mouse line carrying an independent copy of the 10 gRNAs used in the original CARLIN construct, with the goal of enhancing barcode editing. By crossing homozygous mice having all three target arrays with homozygous *Cola1a1*^tetO-Cas9–TdT-gRNA/tetO-Cas9–TdT-gRNA^:*Rosa26*^M2-rtTA/M2-rtTA^ mice, we obtained DARLIN-v1 mice (Figure 3A). Unless otherwise stated, all the data presented below were generated from this DARLIN-v1 line, referred to hereafter simply as DARLIN mice.

**Figure 3.**
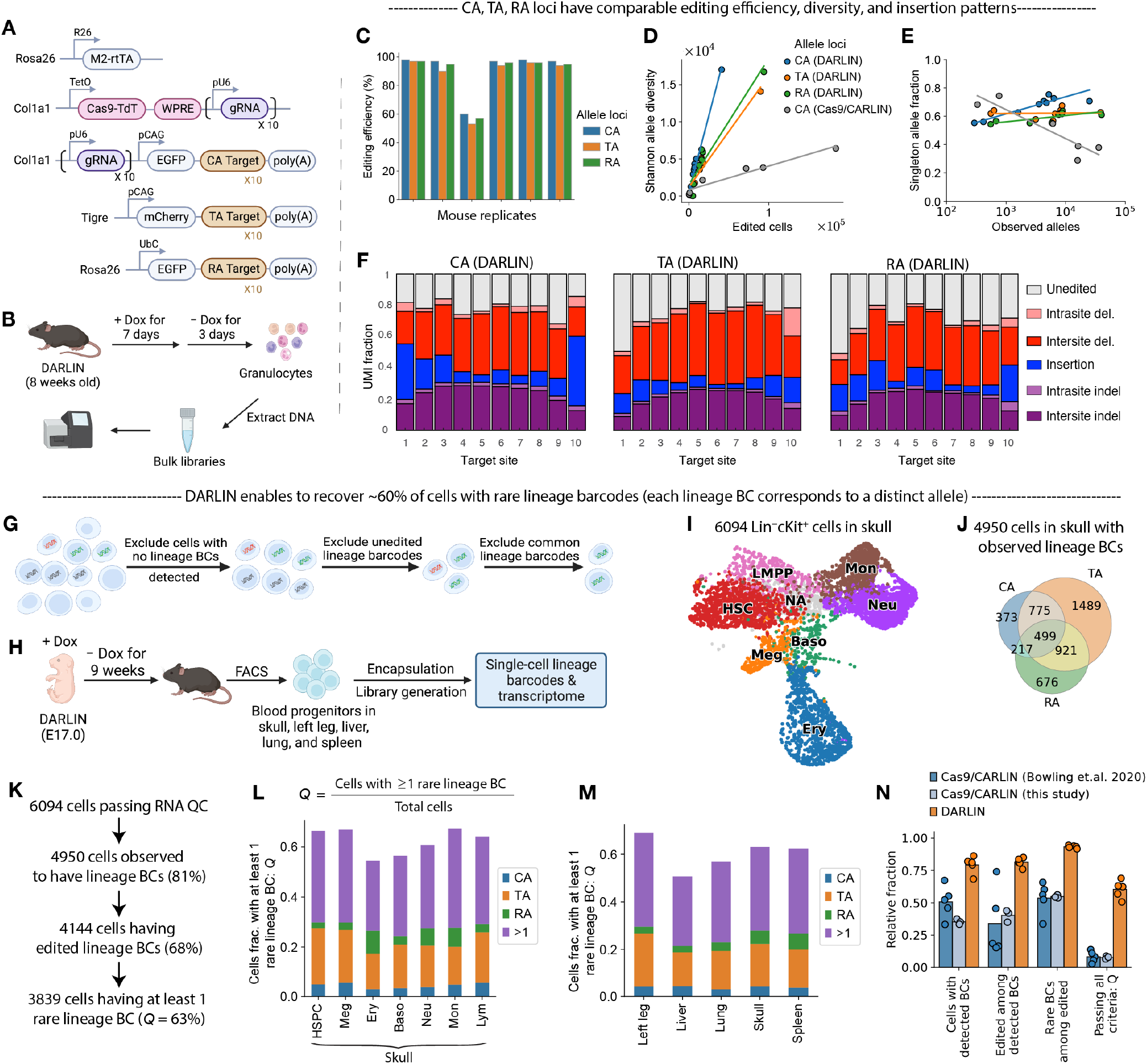
Characterization of DARLIN mice. (A) Genetic elements of the DARLIN system. Compared with DARLIN-v0 (Figure 2A), a second copy of gRNAs was integrated at the *Col1a1* locus after Cas9–TdT. Two additional target arrays bearing the same ten target sites in two different orders were integrated at the *Tigre* and *Rosa26* locus, respectively. (B) Experimental scheme to compare the target array editing in the three loci of DARLIN. (C–F) Analyses of alleles generated from the CA, TA, and RA loci: editing efficiency (C), Shannon allele diversity as a function of the inferred number of edited cells (D), the singleton-allele fraction as a function of the number of observed alleles (E), and frequencies of editing patterns across the ten target sites (F). (G) Schematic of the quality control (QC) pipeline used to select cells for lineage analysis from single-cell data. (H) Experimental scheme by which the single-cell lineage tracing data was generated with DARLIN. (I) UMAP embedding of the transcriptomes measured for the 6,094 Lin^-^cKit^+^ cells from skull-derived bone marrow that passed RNA QC. HSC, hematopoietic stem cell; LMPP, lymphoid-biased multipotent progenitor; Meg, megakaryocyte; Ery, erythrocyte; Baso, basophil; Neu, neutrophil; Mon, monocyte. (J) Venn diagram showing the number of cells in the skull for which each type of target array (CA, TA, or RA) or combination of these was detected. (K) Cell number at each filtering step for the skull dataset. (L & M) Fraction of cells for which a rare allele was detected from at least one target array either for different cell types from the skull (L) or blood cells sampled from different tissues (M). Each bar is colored according to the percentage of cells with alleles at only a single locus (CA, TA, or RA), or multiple loci (>1). (N) Comparison of the single-cell lineage data derived from DARLIN with those of Cas9/CARLIN, in terms of the number of cells that passed each QC step described in G. The DARLIN data are from the adult tissues shown in M, the Cas9/CARLIN data generated in this study were collected from the head, tail, and trunk of a mouse embryo, and the published Cas9/CARLIN data (collected from bone marrow) correspond to those of Figure 6 from Bowling et al., 2020^19^.

To evaluate the editing performance across the CA, TA, and RA loci, we induced six adult DARLIN mice with Dox for 1 week and analyzed the alleles from bone marrow granulocytes with bulk DNA sequencing (Figure 3B). We found that the three target-array loci achieved a similar editing efficiency (Figure 3C), comparable Shannon allele diversity (Figures 3D and S2B), as well as ~60% singleton allele fraction (Figure 3E). We further confirmed that CA, TA, and RA loci had similar editing patterns (Figures 3F and S2C–S2E). These data were highly consistent with the above CA-locus data from the DARLIN-v0 mouse (Figures 2C–2I). We conclude that in the DARLIN mouse line, the TA and RA loci perform comparably to the CA locus with respect to editing efficiency and barcode diversity.

### DARLIN achieves superior single-cell lineage coverage

Because target arrays in DARLIN are transcribed, one can simultaneously profile the lineage barcodes and transcriptomes of single cells. Transcriptomic information enables systematic resolution of cell states, which is crucial for understanding lineage relationships in a heterogeneous population without conventional sorting markers. However, only cells with lineage barcodes that are detected, edited, and rare will be useful for downstream lineage analysis to avoid barcode homoplasy (Figure 3G). We therefore systematically evaluated the characteristics of single-cell lineage tracing in the DARLIN mouse line.

We labeled DARLIN mice at E17.0 and generated a single-cell lineage-tracing dataset of Lin-cKit+ blood cells from skull bone marrow (Figure 3H). We obtained 6,094 cells after quality control (Figure 3I) and detected alleles from at least one target locus in 81% of cells, half of which (overall 40%) contain at least two of three target loci (Figure 3J). The target array from the TA locus had a higher capture rate due to its higher expression (Figures 3J and S2F). Among these 6,094 cells, 3,839 cells (63%) had at least one rare allele (Figure 3K), and this fraction was comparable across the seven blood cell types in the skull (Figure 3L). We profiled hematopoietic cells from four other tissues (left-leg–derived bone marrow, liver, lung, and spleen) in single-cell assays (Figure 3H), and also observed ~60% of cells with at least 1 rare allele (Figures 3M). We classified an allele as rare below a stringent threshold of homoplasy probability, according to a large allele bank (~10^5^ alleles) aggregated across three bulk DARLIN mouse replicates from an experiment designed to measure intrinsic allele frequencies (Figure S4; STAR Methods). We also confirmed that the editing in CA, TA, and RA loci are independent (Figure S2G), implying that the total barcodes in DARLIN could reach ~10^18^ since CA locus alone generates at least 10^6^ alleles. Notably, this barcode complexity far exceeds the total number of cells (~10^10^) in an adult mouse.

Next, we systematically compared the single-cell lineage readout between DARLIN and Cas9/CARLIN^19^ mouse lines. On average, we detected expression from at least one lineage locus in 80% of cells derived from the DARLIN mouse, compared to ~50% in Cas9/CARLIN (Figure 3N). Among the detected lineage barcodes, ~80% were further edited in DARLIN, compared to ~35% in Cas9/CARLIN, which was consistent with our observations of editing efficiency in mouse embryos (Figure S2H) and adult mice (Figure 3C). Finally, among the edited cells, ~93% of cells from DARLIN mice had at least one rare allele, while this fraction was only 55% for Cas9/CARLIN mice. In total, DARLIN mice achieved ~60% of cells passing all three filters (i.e., had an allele that was detected, edited, and rare), compared with ~10% in Cas9/CARLIN. Our assessment of Cas9/CARLIN agreed with our previous assessment^19^, suggesting a similar stringency of our approach for identifying rare alleles (which may be relaxed when labeling fewer cells), and was consistent across two datasets (Figure 3N). Together, the above data demonstrate that the DARLIN mouse line has superior single-cell lineage coverage and a barcode diversity that exceeds the number of cells in an entire adult animal.

### Mapping cell fate choices among unperturbed blood progenitors in vivo

We next demonstrated the utility of DARLIN to study cell fate choice during developmental hematopoiesis. Several studies have shown that hematopoietic stem and progenitor cells (HSPCs) can be divided into subpopulations with functional heterogeneity^10,21,22,41^, including subsets with distinct fate biases. However, when this fate bias is established during development, and the molecular features of these biased HSPCs, remain poorly understood.

We re-analyzed our single-cell lineage tracing data from skull-derived bone marrow induced at E17.0 and collected in adulthood (Figure 3H). We identified six major cell types among the 6,094 profiled single cells: hematopoietic stem cells (HSCs), lymphoid-biased multipotent progenitors (LMPPs), megakaryocyte progenitors (Meg), erythrocytes, neutrophils, and monocytes (Figures 3I, 4A, and S4A). We integrated information from the three target-array loci to assign a clone ID to each cell (STAR Methods). In total we identified 948 distinct clones (Figure 4B): some clones occupied multiple cell fates (Figure 4C, left panel), while others had only one observed fate outcome (Figure 4C, middle and right panel). With this data, we asked if some HSPCs (HSCs and LMPPs) demonstrated differentiation bias towards specific fates (Figure 4A).

**Figure 4.**
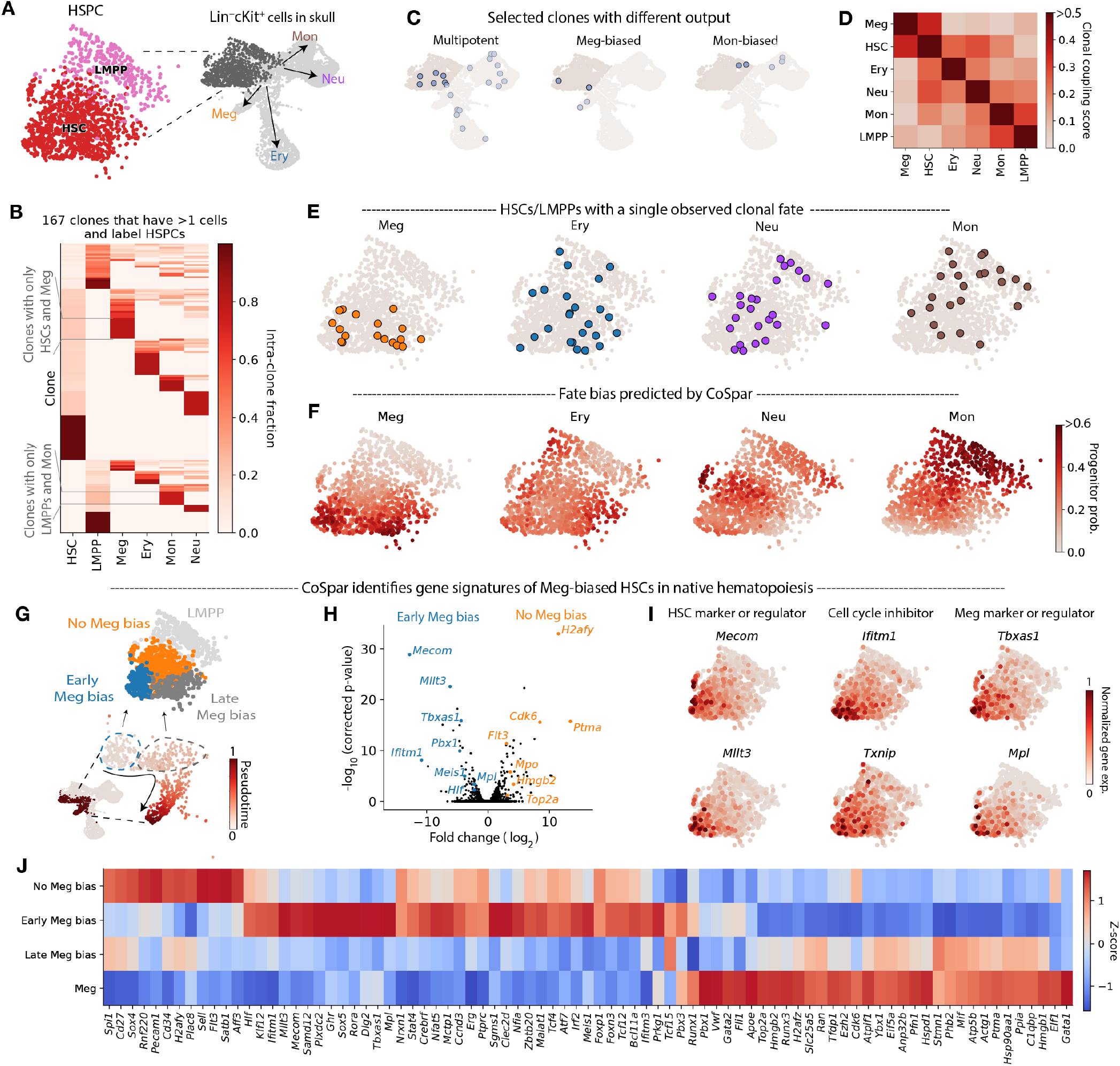
Early Fate Priming Among Hematopoietic Stem and Progenitor Cells (HSPC) (A) UMAP embedding of HSPCs (HSCs and LMPPs) from Lin^-^cKit^+^ cells derived from skull bone marrow (See also Figure 3I). (B) Clonal profile of the normalized proportion of each annotated cell type (column) within each clone (rows). Only the 167 clones labeling HSPCs are shown. (C) Selected clones with multipotent fate outcomes, or a single observed clonal fate. (D) Heatmap of clonal coupling scores across major cell types (STAR Methods). (E) UMAP embedding of HSPCs redrawn from (A), labeled to indicate HSPCs that were clonally associated with a single mature cell type (Meg, Ery, Neu, or Mon). (F) CoSpar-predicted probability of each HSPC to generate each of the four observed mature cell types (Meg, Ery, Neu, or Mon). (G) Identification of early Meg-biased HSCs using the inferred differentiation trajectory from HSCs into Meg calculated by CoSpar. (H) Volcano plot of differentially expressed genes when comparing early Meg-biased HSCs with inferred HSCs without Meg biases. (I) Expression of selected genes on top of the UMAP embedding for HSPCs. (J) Heatmap showing the expression of selected genes across different HSPC clusters and Meg. Z-scores were calculated from the mean and standard deviation of each gene’s expression in the four cell populations.

The clonal coupling scores across major cell types (i.e., a normalized correlation to measure how often two cell types jointly appear within the same clone; STAR Methods) suggested a strong lineage coupling between Meg and HSC (but not between Meg and other cell types), and between monocyte and LMPP (Figure 4C and Figure S4B). This agrees with earlier reports that a subset of HSCs can directly generate Meg^8,22,42–45^, and that LMPPs are primed to generate monocytes rather than neutrophils^21^. Considering that barcoding was induced at E17.0, a developmental time point when HSCs still reside in the fetal liver, our data suggest that HSCs at this time already carry functional features that will be evident even after their migration to the bone marrow. Thus, Meg bias is likely to arise earlier than what has been previously reported^46^. We found that 48% of our 175 clones that labeled both HSPCs and a later fate had a single clonal fate (Figure 4D), suggesting the possibility of early fate bias. Inspecting those HSPCs that are clonally associated with a single mature fate, we found that only Meg-biased clones had distinct transcriptomic signatures (Figure 4E). We previously developed CoSpar, a computational approach that utilizes coherent and sparse lineage dynamics to robustly infer early cell fate choice^47^. We applied CoSpar to infer early fate priming by integrating transcriptomic and lineage information. Consistent with the above observations, CoSpar predicted that Meg originate specifically from a subset of HSCs (Figure 4F, left panel). Interestingly, CoSpar also predicted that monocytes originate predominantly from LMPPs (Figure 4F, right panel), which agreed with our clonal coupling analysis (Figure 4C). Importantly, we failed to infer such early fate bias when down-sampling our DARLIN data to match the frequency of cells with detected, edited, and rare alleles in Cas9/CARLIN data (Figure S4D).

Next, to identify the early transcriptomic signature of Meg-biased HSCs, we inferred the differentiation trajectory from HSCs to Meg using the above CoSpar predictions, then split the Meg-biased HSCs into two populations based on their pseudotime: early or late Meg bias (Figure 4G). Compared with HSCs without Meg bias, the early Meg-biased HSCs exhibited enriched expression of genes involved in maintaining long-term HSC identity (*Mecom, Mllt3*, and *Hlf*), cell-cycle inhibition (*Ifitm1, Txnip*, and *Ifitm3*), and megakaryopoiesis regulation (*Tbxas1*, *Mpl*, and *Meis1*) (Figures 4H and 4I)^22^. We also identified many genes without an established association with Meg bias in HSCs (Figure 4J), including the transcription factors *Klf12, Sox5, Rora, Pbx3, Pbx1*, and *Gata2* (Figure S4C). Taken together, our analyses demonstrated that the DARLIN mouse line generates high-quality single-cell lineage tracing data that resolves early fate priming within HSCs, leading to the identification of novel gene signatures of Meg-biased HSCs in unperturbed hematopoiesis *in vivo*.

### Lineage relationships of blood cells across bones reveal HSC migration dynamics over development and adulthood

Next, we used the DARLIN mouse line to systematically evaluate clonal dynamics of the migration of hematopoietic progenitors over development and adulthood (Figure 5A). In murine hematopoiesis, definitive blood progenitors arise at E10.5 with the formation of *Runx1*-expressing clusters within the aorta-gonad-mesonephros (AGM) region in the embryo^48^. At around E11.5, these progenitors migrate to the fetal liver where they undergo rapid expansion before colonizing the bone marrow at around the time of birth. The clonality of bone-marrow colonization and the extent of HSPC circulation during ontogeny are particularly unclear. The process of HSC circulation has previously been studied in the adult mouse by parabiosis^49–51^. Although Wright *et.al.* observed that up to 8% of HSCs migrated from one mouse to the other over 39 weeks^49^, a later study only observed 1–2.5% migratory HSCs^50^. Furthermore, parabiosis experiments often lead to injury and inflammation, which might influence the behavior of HSPCs in such studies. The high barcoding capacity of the DARLIN model presented us with the unique opportunity to address these questions at the level of individual clones in a completely physiological context.

**Figure 5.**
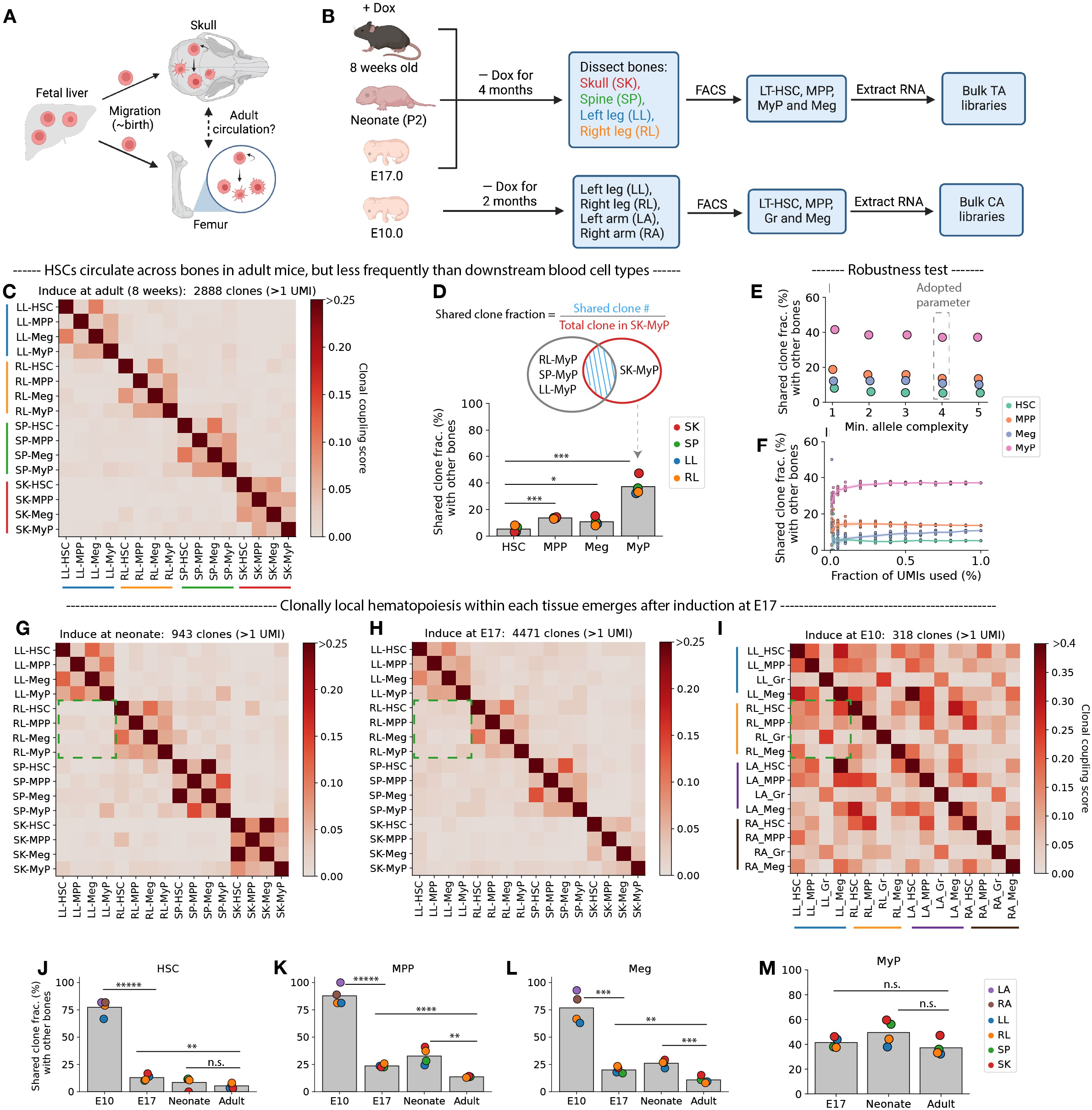
Lineage Relationships of Blood Cells across Bones. (A) Schematic of migration of blood progenitors across developmental stages. (B) Experimental design to investigate the extent of HSC migration and inter-bone circulation using the lineage relationships of blood cells within and between bones.. Leg bones include femur, tibia and fibula, while arm bones include humerus, ulna and radius. (C–F) Clonal analysis of bulk lineage-tracing data from week-8 induction. (C) Heatmap of clonal coupling scores between cell types from each bone. (D) Shared clone fraction of each cell type across bones (\**p* < 0.05; \*\**p* < 0.01; \*\*\**p* < 0.001; \*\*\*\**p* < 10^-4^; \*\*\*\*\**p* < 10^-5^, *t*-test). (E) Shared clone fraction for each cell type averaged over all four bones calculated over a range of thresholds for excluding low-complexity (and therefore likely more common) alleles. (F) Shared clone fraction when performing different amounts of UMI down-sampling. (G–I) Heatmap of clonal coupling scores between cell types across bones for mice induced at the neonate stage (G), E17.0 (H), and E10.0 (I). (J–M) Shared clone fraction with other bones when editing was induced at different developmental stages for HSCs (J), MPP (K), Meg (L), and MyP (M). Each point is colored by the bone for which the shared clone fraction was calculated.

We induced DARLIN mice at different developmental stages (adulthood, neonate, and E17.0). After four months, we dissected bone marrow from four locations (skull, spine, left leg (i.e., femur, tibia, and fibula), and right leg), and used FACS to sort long-term (LT) HSCs, MPPs, myeloid progenitors (MyPs), and Meg from each bone to profile their lineage barcodes via bulk RNA-sequencing (Figure 5B, upper panel, and S5A). In a separate experiment, we also induced one DARLIN mouse at E10.0, waited 2 months, and profiled the lineage barcodes across major blood cell types sorted from four different bones (Figure 5B, lower panel).

First, we determined to what extent hematopoietic progenitors circulate between different bones during adulthood by looking at the mouse induced in adulthood. The presence of a clone in more than one bone indicated inter-bone migration, by which an individual HSC (or progenitor) divides and colonizes a different bone marrow niche. Calculating the clonal coupling scores between all pairs of cell types from all sorted populations, which accounts for clone identities and their sizes, we observed that hematopoietic populations were strongly related in clonal origin within each bone, but not between bones (Figure 5C). This is consistent with the idea that hematopoiesis is predominantly maintained locally within each bone in the adult, at least within four months. Indeed, each of the clones resided predominantly in one bone, with only a small fraction of cells (UMIs) detected in other bones (Figure S5A). Considering the fraction of HSC-containing clones found in one bone (e.g., skull HSCs) that are also detected in HSCs from other bones (irrespective of clone size), we observed ~5% of HSC-containing clones were shared with HSCs from at least one other bone (Figure 5D). The overlap fraction increased significantly to ~14% for MPPs (*t*-test, *p* < 10^-3^), and to ~40% for MyPs (*p* < 10^-3^) (Figure 5D). To exclude contamination from common alleles, we only used *de novo* alleles (~80% of all alleles) from this experiment that were not found in our pre-assembled allele bank with ~100,000 alleles (Figure S5C). Accordingly, the inferred shared clone fraction was robust to i) the mutational complexity of the alleles considered (Figure 5E), ii) down-sampling UMIs (Figure 5F), and iii) read cutoffs used for allele calling (Figure S5D). Our data extend the earlier findings of HSC circulation between bones^49,50^, and provide evidence that this process actively occurs in a native physiological context. Our findings also demonstrate that more differentiated populations like MPPs and MyPs circulate more actively.

We next studied the dynamics of inter-bone migration in the neonate. It is unclear whether the migration of post-birth HSCs, which likely have colonized bone marrow, is more dynamic than those in the adult, although indirect evidence suggests that this might be the case^52^. Our results demonstrated that at four months after birth, local hematopoiesis was still prevalent even when barcoding was induced in the neonatal stage (Figure 5G). There was ~8% shared clone fraction of HSCs between the bones studied (Figure S5E and S5F), similar to induction in adulthood (*p* = 0.34). Thus, our results argue against a significantly increased rate of inter-bone migration after birth in comparison to later in adulthood.

We also induced at E17.0, a stage when HSCs are predominantly located in the fetal liver^53^. Remarkably, we still observed local hematopoiesis within most clones four months later (Figures 5H and S5A). On the other hand, when a mouse was labeled at E10.0, the clonal coupling scores of blood cells within the same bone were comparable to those between different bones (Figure 5I), and an ~80% shared clone fraction between bones was observed for different cell types (Figure S5E and S5F). This was expected, because at E10.0 lineage labeling occurs before the rapid HSC expansion in the fetal liver and subsequent migration to the bone marrow, leading to initial colonization of multiple bones by the same clone. Thus, our observations suggest that HSCs labeled in the late fetal liver stages (E17.0) predominantly seed one single bone microenvironment, likely due to limited clonal expansion before colonizing the bone marrow^53^. The shared clone fractions from induction at E17.0 were still higher than those of induction at adulthood for HSCs, MPPs, and Meg (Figures 5J–5L), implying that some clonal expansion before migration may contribute to the relatively higher shared clone fractions of E17.0-labeled hematopoiesis. We observed that MyPs had comparable shared clone fractions when labeling across these stages (Figure 5M). This difference was consistent with MyPs undergoing more active circulation in the adult stage. In conclusion, the high barcoding capacity of the DARLIN model has allowed us to obtain unique insight into the process of HSC migration during development and adulthood.

### Camellia-seq simultaneously profiles chromatin accessibility, DNA methylation, gene expression, and lineage information in single cells

Integrating lineage tracing with single-cell transcriptomic measurement enables systematical dissection of fate biases for a transcriptomically heterogeneous population^20–22,47^. The epigenetic state of a cell also plays a crucial role in regulating its dynamics and function^23–25^. An integrative measurement of lineage, transcriptome, and epigenome at the single-cell level would enable a deeper understanding of how cell fate choice is regulated and how cell identity is maintained across different modalities. Here, we developed a sequencing method to simultaneously measure <u>c</u>hromatin <u>a</u>ccessibility, DNA <u>me</u>thylation, gene expression, and <u>li</u>ne<u>a</u>ge information in single cells (Camellia-seq) (Figure 6A). Camellia-seq extends scNMT-seq^54–57^ by incorporating lineage barcode measurement. Briefly, a single cell is split into nuclear and cytoplasmic fractions. Endogenous mRNAs and expressed lineage barcode transcripts are reverse-transcribed and amplified from the cytoplasmic fraction via a modified STRT-seq protocol^58,59^. The nuclear fraction is treated with GpC methyltransferase, which preferentially methylates cytosine from GpC dinucleotides within regions of open chromatin^60^. The endogenous DNA methylation (methylated cytosine in CpG dinucleotides) and accessible chromatin (methylated cytosine in GpC dinucleotides) are then profiled with single-cell Bisulfite Sequencing^61^.

**Figure 6.**
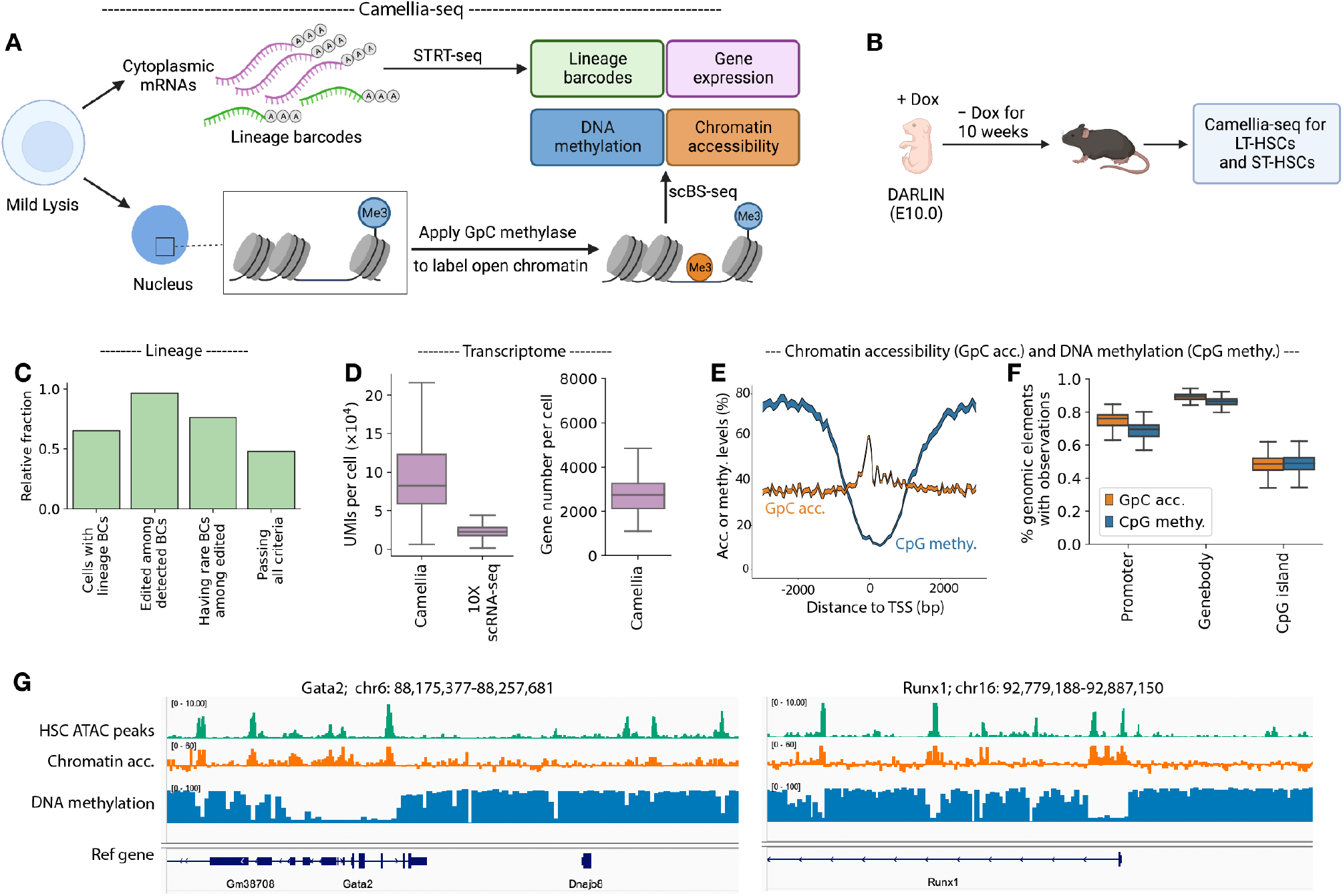
Joint Profiling of Lineage, Gene Expression, Chromatin Accessibility, and DNA methylation with Camellia-seq. (A) Schematics of Camellia-seq to profile lineage barcodes, gene expression, DNA methylation, and chromatin accessibility in single cells. (B) Experimental scheme to profile bone-marrow LT- and ST-HSCs with Camellia-seq. (C) Fraction of cells that passed each QC step described in Figure 3G. (D) Box plots showing the number of observed UMIs per cell (left) and the number of expressed genes per cell (right) for the scRNA-seq data generated with Camellia-seq. The number of observed UMIs per cell for the scRNA-seq data generated with the 10X genomics protocol (Figure 3I) is also shown. (E) The average chromatin-accessibility or DNA-methylation profile over the transcription start sites (TSS) of ~20,000 different genes in a cell. (F) Box plot showing the genomic coverage across promoters, gene bodies, and CpG islands. Each GpC site must be covered by ≥3 reads, and CpG site by ≥1 read. (G) Pseudobulk chromatin accessibility and DNA methylation surrounding the TSS of *Gata2* and *Runx1*. The bulk HSC ATAC-seq peaks from Li et al., 2020 are also shown.^53^

We profiled HSCs with Camellia-seq to evaluate the quality of each data modality. We induced lineage labeling in the DARLIN mouse at E10.0 when HSCs just form in the AGM region, waited nine weeks until adulthood, and extracted both long-term HSCs (LT-HSCs) and short-term HSCs (ST-HSCs) from adult bone marrow to perform Camellia-seq (Figure 6B). Approximately 50% of the single cells profiled with Camellia-seq had a rare lineage barcode (Figure 6C). We observed a median transcriptomic abundance of ~100,000 UMIs derived from ~3,000 genes per cell (Figure 6D). Using epigenomic modalities from single cells, we reproduced the stereotypic pattern that DNA methylation drops within 1-kb of the transcription starting site (TSS), while the chromatin accessibility peaks near the TSS and drops with an oscillatory pattern in the direction of transcription initiation^54^ (Figure 6E). Furthermore, Camellia-seq achieved a high genomic coverage: ~70% of promoters and ~90% of the gene bodies were represented with at least 3 detected GpC sites and 1 CpG site (Figure 6F). By aggregating single-cell epigenomic measurements into a pseudo bulk dataset, we further confirmed that the resulting chromatin accessibility measurements agreed with bulk ATAC-seq measurements of HSCs from a published dataset^53^ and anti-correlates with DNA methylation measurements in our data (Figure 6G).

## Discussion

Here, we describe DARLIN, a lineage-tracing mouse line with a superior lineage barcoding capacity and enhanced single-cell lineage coverage. Building on the current Cas9/CARLIN lineage-tracing system, we first incorporated TdT to increase insertion events in lineage barcodes, and subsequently expanded DARLIN to include three independent lineage-recording loci. DARLIN can theoretically generate an estimated 10^18^ unique lineage barcodes, and has ~90% barcode editing efficiency compared to ~30% in Cas9/CARLIN (Figures 2J, 3C, 4N, and S2H). In single-cell assays, ~60% of captured cells from DARLIN have edited and rare lineage barcodes (due to reduced barcode homoplasy) for further clonal analysis, compared to ~10% in the Cas9/CARLIN mouse line. This translates to more useful clones per sample (due to higher editing efficiency and barcode diversity), more cells per clone (due to higher lineage barcode capture efficiency per cell), and dramatically reduced experimental costs for generating a dataset with sufficient clonal information to address a biological question. Finally, like the Cas9/CARLIN mouse line, the lineage barcoding in DARLIN can be induced at any time point and across a wide range of tissues, and DARLIN is a stable and genetically defined mouse line that can be shared across a wide biological community.

The large barcode diversity is useful in two ways. First, a system with up to 10^18^ barcodes is in principle sufficient to uniquely label every one of the ~10^10^ cells of an adult mouse. Thus, the DARLIN mouse line can be used to study large biological systems, such as adult tissue homeostasis (as we demonstrated for hematopoiesis in this study), inflammation response, and tissue injury and repair. Secondly, noting that barcodes occur at different frequencies, a larger barcode diversity results in a higher fraction of rare clones in our data (~93% of edited cells having rare alleles in Figure 3N). In many applications, profiling alleles from a single target array locus may already provide sufficient lineage information. Having secondary and tertiary sets of independent barcodes in the same samples will increase the robustness of lineage measurements.

In our applications, we demonstrated first that the DARLIN mouse line enables the study of early fate bias within native HSCs at high resolution, leading to the identification of multiple new genes correlated with Meg bias in HSCs. In our second application, we studied the lineage relationship of hematopoietic cells across different bones. Our data provide the first demonstration of HSC inter-bone migration in a completely physiological context, with a ~5% shared clone fraction of HSCs between different bones accumulated over four months after induction in adulthood. These observations support the idea that HSCs continuously circulate at low levels in adulthood^49^. Considering that we dissected only ~70% of the mouse bone marrow, we likely underestimated the extent of HSC migration. We predict that the ~5% shared clone fractions measured for HSCs would increase with a longer waiting time after barcode induction. Our experiments also provide insight into migration from the fetal liver and during early postnatal growth. We observed comparable HSC migration between barcode induction in the neonatal and adult stage. Even for induction at E17.0, when HSCs only begin to migrate from the fetal liver to bone marrow, we observed within-bone or local hematopoiesis across most clones detected four months later. Our findings at this stage confirm our observations of limited migration post initial bone settlement.

In parallel, we have established Camellia-seq to simultaneously profile lineage barcodes, chromatin accessibility, DNA methylation, and gene expression in single cells. Using DARLIN, we showed that Camellia-seq generates high-quality data for each of the modalities.

Additionally, Camellia-seq is compatible with any lineage-tracing approach where the lineage barcodes are transcribed as mRNA^31^. DARLIN mice may also be induced with Dox over a series of time points to generate alleles with a hierarchical structure to obtain more hierarchical cellular lineage relationships of large numbers of cells during tissue development or homeostasis. The lineage barcodes in DARLIN may also be resolved spatially to understand spatial lineage dynamics in tissues. Overall, the DARLIN mouse line and Camellia-seq method provide a powerful tool for studying the relationships and underlying molecular mechanisms of diverse biological processes.

## Limitations

The target arrays in DARLIN still suffer from array deletions, which might limit their use for lineage reconstruction across multiple cell divisions. This may be circumvented by generating sequential mutation events along the recording array to produce hierarchically labeled clones^62^.

Camellia-seq is currently a low-throughput and costly plate-based method that requires deep genomic coverage for each cell. A cost-effective and high-throughput method would be desirable.

## ACKNOWLEDGMENTS

We are grateful to members of F.D.C laboratory; and BCH Mouse Gene Manipulation Core for ESC injection and chimera generation; and Ronald Mathieu, Mahnaz Paktinat, Ranjan Maskey and Betelhem Gemechu from HSCI-BCH Flow Cytometry Research Lab for guidance and assistance with FACS; Alejo Rodriguez-Fraticelli for assistance with designing the Cas9–TdT plasmid; Wei-Chien Yuan for assistance with hydrodynamic injections in mice; Fan Zhou and Chloé Baron for assistance with mouse AGM dissection and pre-HSC sorting; Fuchou Tang and Shuhui Bian for assistance with multi-omic experiments and analysis. We thank Qiu Wu from the Klein lab for constructive comments on our manuscript. S.-W.W. is supported by the Damon Runyon Quantitative Biology Award from the Damon Runyon Cancer Research Foundation (02-20). Illustrative figures are created with BioRender.

## AUTHOR CONTRIBUTIONS

L.L., S.-W.W. and F.D.C. conceived the project, designed the study, and analyzed the data. L.L., S.-W.W. and F.D.C. wrote the manuscript with help from all other authors. L.L. generated Cas9–TdT, Cas9–TdT-gRNAs and Cas9–TdT-gRNAs-TA mESCs, and Cas9–TdT, Cas9–TdT-gRNAs mouse lines. S.B. and B.L. produced Tigre-target-array and Rosa26-target-array mESCs and mouse lines. L.L. bred DARLIN-v0 and DARLIN mouse lines and developed the Camellia-seq method. L.L. performed the mESCs, animal and sequencing experiments with help from Q.Y. (sample collection and cell sorting), K.A. (mouse perfusion and liver dissociation), S.B. (mouse AGM dissection) and M.F. (IP injection in mouse neonates). S.-W.W. developed the computational methods for analyzing DARLIN and Camellia-seq data, wrote the software packages, and performed all the statistical analyses. S.E.M. provided valuable feedback on data analysis and interpretation. A.M.K. supervised statistical analysis and lineage-tracing experimental design. F.D.C. and S.-W.W. supervised the study.

